# Labor- and cost-effective long-read amplicon sequencing using a plasmid analysis service: Application to transposon-inserted alleles in Japanese morning glory

**DOI:** 10.1101/2024.09.19.613814

**Authors:** Soya Nakagawa, Atsushi Hoshino, Kyeung-Il Park

## Abstract

The sequencing of PCR fragments amplified from specific regions of genomes is a fundamental technique in molecular genetics. Sanger sequencing is commonly used for this analysis; however, amplicon sequencing utilizing next-generation sequencing has become widespread. In addition, long-read amplicon sequencing, using Nanopore or PacBio sequencers to analyze long PCR fragments, has emerged, although it is often more expensive than Sanger sequencing.

Recently, low-cost commercial services for full-length plasmid DNA sequencing using Nanopore sequencers have been launched in several countries, including Japan. This study explored the potential of these services to sequence long PCR fragments without the need for cloning into plasmid DNA. PCR fragments of 4–11 kb, amplified from the *DFR-B* gene involved in the biosynthesis of anthocyanin, with or without *Tpn1* transposons in Japanese morning glory (*Ipomoea nil*), were circularized using T4 ligase and analyzed as templates. Although some inaccuracies in the length of homopolymer stretches were observed, the remaining sequences were obtained without significant errors. This method could potentially reduce the labor and costs associated with cloning, primer synthesis, and sequence assembly, thus making it a viable option for the analysis of long PCR fragment sequences. Moreover, this study reconfirmed that *Tpn1* transposons are major mutagens in *I. nil* and demonstrated their transposition in the Violet line, a long-used standard in plant physiology.

## Introduction

Sequencing of genome fragments amplified by PCR is routinely performed in laboratories worldwide. Although Sanger sequencing has traditionally been the standard method used, amplicon sequencing using next-generation sequencing (NGS) has become increasingly common, particularly for microbiome analysis (Caporaso et al. 2011). Moreover, long-read amplicon sequencing, which can sequence PCR products greater than 10 kb in length using technologies such as PacBio HiFi and Oxford Nanopore Technologies sequencing, has recently emerged (Karst et al. 2021). However, to sequence a single PCR product, the NGS throughput is excessive. Consequently, multiplex analysis, which allows for the simultaneous sequencing of many PCR products, is often employed to reduce the cost per sample. Nonetheless, Sanger sequencing remains the most common approach when analyzing a small number of samples.

Although long-read amplicon sequencing services are available, they tend to be more expensive than Sanger sequencing. However, low-cost full-length plasmid DNA sequencing services using Nanopore technology have recently been launched by several companies (e.g., Eurofins Genomics (USA), AZENTA Life Sciences (USA), and Plasmidsaurus (USA)) and are currently available in Japan. Utilizing these services can substantially reduce the labor, cost, and time required compared to Sanger sequencing through primer walking. Moreover, this reduction increases with the analysis of longer DNA sequences. In this study, a commercial plasmid DNA sequencing service was used to determine the sequence of circularized PCR products, which were created to resemble plasmid DNA through ligation (Fig. 1).

**Figure 1.**
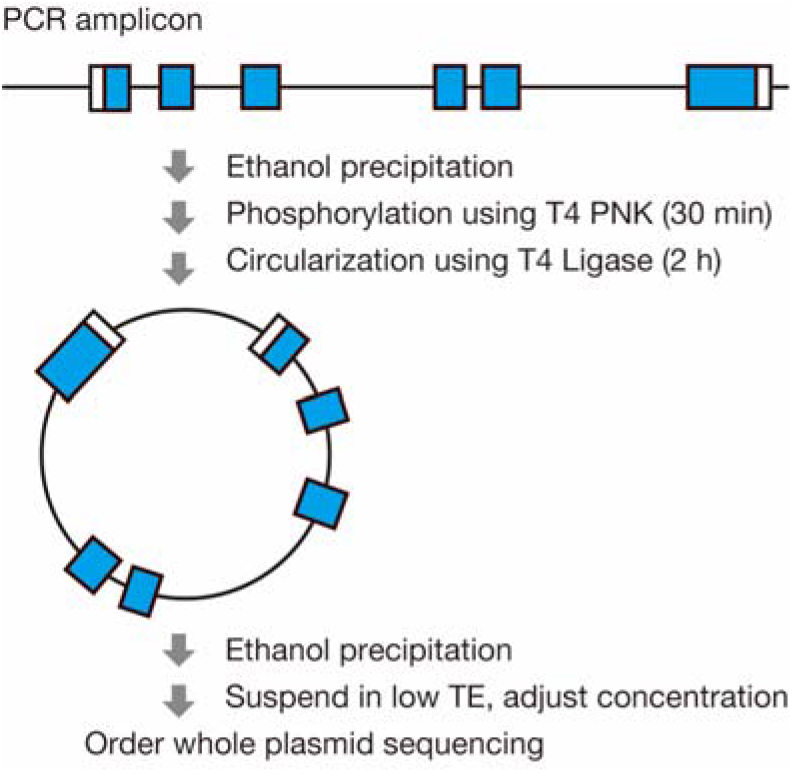
Flowchart for the preparation of templates from PCR amplicons for whole plasmid DNA sequencing services using Nanopore sequencers. T4PNK; T4 Polynucleotide Kinase.

Japanese morning glory, *Ipomoea nil*, is a unique bioresource of Japan. In domestic lines, the *Tpn1* family transposons exhibit high transposition activity, which leads to numerous insertion mutations (Inagaki et al. 1994; Fukada-Tanaka et al. 2000; Hoshino et al. 2001; Nitasaka 2003; Iwasaki and Nitasaka 2006; Hoshino et al. 2009; Morita et al. 2014; Morita et al. 2015; Hoshino et al. 2016). Most of these mutations affect the flower color and patterns as well as the morphology of flowers and leaves, due to selective breeding for horticultural purposes. In the Tokyo Kokei Standard (TKS) line, the whole-genome sequence revealed 340 copies of the *Tpn1* transposons (Hoshino et al. 2016). The majority are non-autonomous transposons that lack transposase genes, with an average length of 7 kb. Their internal sequences contain fragments of multiple host genes, thereby contributing to their diversification (Takahashi et al. 1999; Kawasaki and Nitasaka 2004). Subterminal repetitive regions (SRRs) consisting of 104- and 122-bp tandem repeats are located at the 5′ and 3′ ends, respectively (Hoshino et al. 1995; Kawasaki and Nitasaka 2004). The length of the terminal regions, including SRRs, approaches the maximum read capacity of capillary sequencers, making it challenging to determine full-length sequences. Of the eleven *Tpn1* transposons identified as causing mutations, the full-length sequences of five have not yet been determined (Hoshino et al. 1995; Nitasaka 2003; Iwasaki and Nitasaka 2006; Hoshino et al. 2009; Morita et al. 2015). This study analyzed the full-length sequences of three novel *Tpn1* transposons that confer white flowers by inserting them into the *DFR-B* gene, which participates in anthocyanin biosynthesis (Fig. 2), using a commercial service for full-length plasmid DNA sequencing.

**Figure 2.**
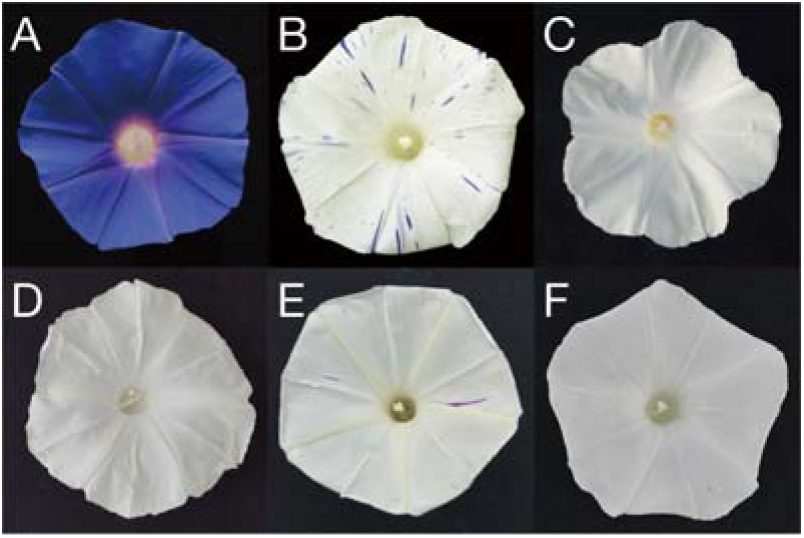
Flower phenotypes: (A) TKS, (B) AK006, (C) AK007, (D) AK010, (E) AK127, and (F) AK205.

## Results and Discussion

As shown in Fig. 1, the analysis of PCR products circularized with T4 ligase using a full-length plasmid DNA sequencing service was validated with two known sequences. First, the sequence of the *DFR-B* gene region of the TKS line (Fig. 2A), whose whole genome has previously been sequenced, was determined (Hoshino et al. 2016). A total of 1,608 reads with an average length of 1,930 bp were obtained, and 1,585 of these reads were used to produce a 4,130 bp circular sequence (Supplementary Table S1, S2). From this circular sequence, a linear sequence was constructed with the PCR primer sequences at both ends. This sequence was identical to the published TKS genome sequence, except for a 1 bp deletion at a 12-bp polythymine stretch in intron 1 (Fig. 3A). Next, the *DFR-B* gene sequence of AK006 (Fig. 2B) was analyzed in the same manner as that of TKS, which resulted in a 10,544 bp sequence. This line harbors the *a3-flecked* allele, in which *Tpn1* is inserted into the *DFR-B* gene (Fig. 3A, B). Comparison of this allele sequence, which was previously determined in AK009, also known as KK/SSB-4 (Hoshino et al. 1995; Takahashi et al. 1999), with the newly obtained sequence revealed a 1 bp deletion in intron 1, identical to the result for TKS (Fig. 3A), along with 1–2 bp deletions at three positions within the *Tpn1* sequence in the 11–15 bp polythymine stretches (Supplementary Fig. S1). These results indicate that the accuracy of this long-amplicon sequencing is approximately 99.9%, with sequencing errors tending to occur in homopolymer stretches. These results are in agreement with a previous report that homopolymeric regions or regions with short repeats account for approximately half of all sequencing errors in Nanopore sequencing (Delahaye and Nicolas 2021).

**Figure 3.**
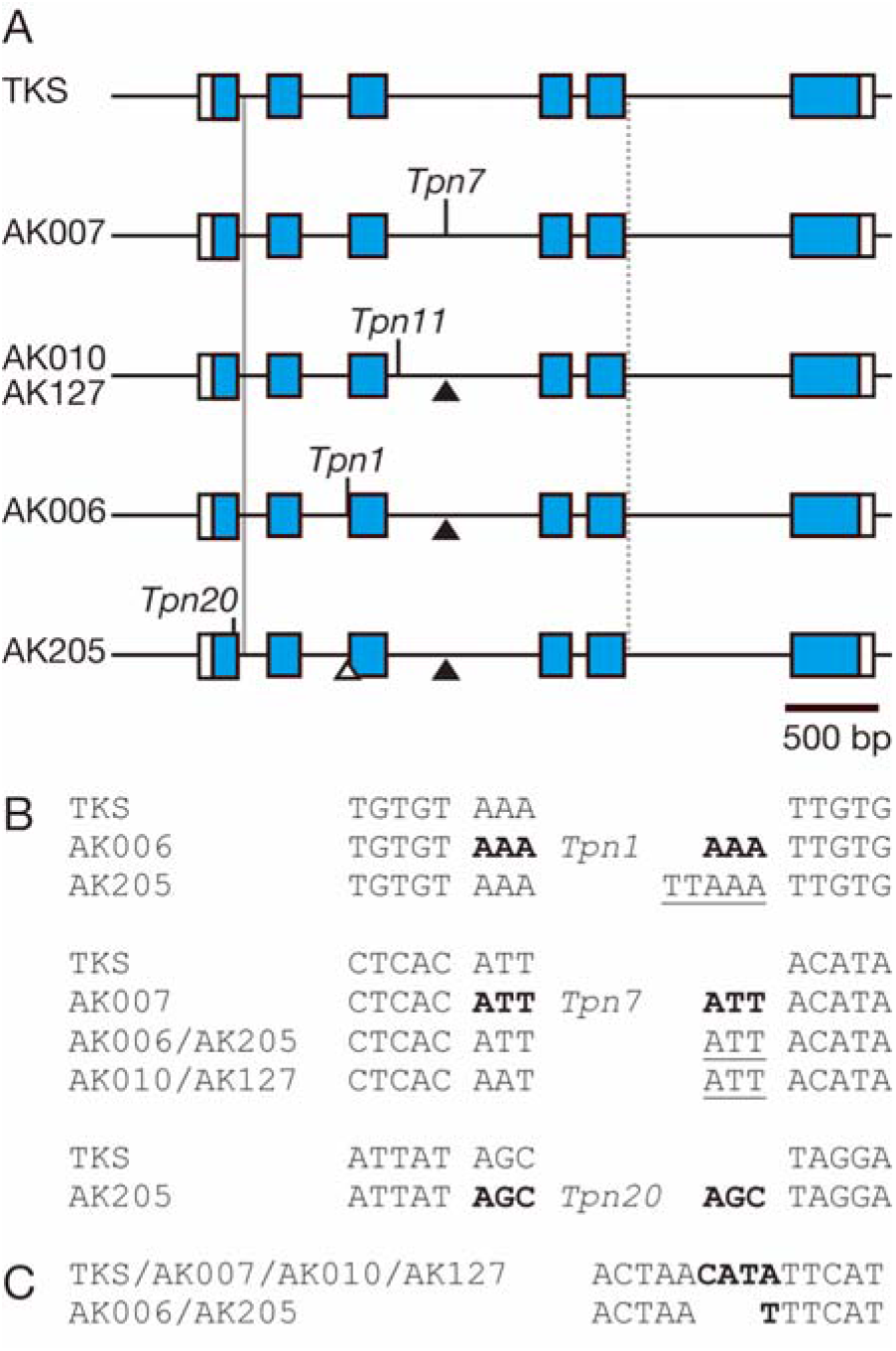
Characterization of the *DFR-B* gene harboring the *Tpn1* transposons. (A) Structure of the sequenced PCR fragments containing the *DFR-B* genes with *Tpn1* transposon insertions. The boxes indicate the exons, with the coding regions shaded in blue. Vertical black bars indicate the transposon insertion sites. White and black arrowheads mark the putative footprints of *Tpn1* and *Tpn7*, respectively. Vertical gray and dotted lines indicate the locations of the 12-bp polythymine stretch and the polymorphic sequences shown in (C), respectively. (B) Sequences of the *Tpn1* transposon insertion sites and their putative footprint sequences. Target site duplications are indicated in bold, and the footprint sequences are underlined. (C) The polymorphic sequences in intron 5 are highlighted in bold.

The long-read amplicon sequencing method was applied to the uncharacterized *DFR-B* alleles with novel *Tpn1* transposons. In AK007 (Fig. 2C), a 6,776 bp *Tpn1* transposon, named *Tpn7*, was identified in intron 3 (Fig. 3A, B). Despite being independently isolated, AK010 and AK127 (Fig. 2D, E) possessed the same allele with a *Tpn1* transposon inserted in intron 3 (Fig. 3A, B); this transposon was designated *Tpn11*. Based on the origins of AK010 and AK127, it is speculated that seeds sold by Marutane Co., Ltd. for the Violet line may have included seeds derived from chimeric individuals containing the *DFR-B::Tpn11* allele. Furthermore, this indicates that *Tpn1* transposons are active in the Violet line, a long-used standard in plant physiology. In AK205 (Fig. 2F), another *Tpn1* transposon insertion was found in exon 1, and named *Tpn20* (Fig. 3A, B). AK205 originated from a germinal revertant of the *DFR-B::Tpn1* (*a3-flecked*) allele, as AK205 and AK006 are descendants of K37. A 5-bp insertion, presumed to be a *Tpn1* footprint sequence, was detected (Fig. 3B). In addition, 3-bp insertion sequences, likely the *Tpn7* footprint, were found in all of the analyzed *DFR-B* mutant lines, which suggests that these lines share a common ancestor plant harboring the

*DFR-B::Tpn7* allele that bears variegated flowers (Fig. 3B). Aside from the insertion of the *Tpn1* transposon, a polymorphism was found in intron 5 (Fig. 3C). In AK006 and AK205, a single thymine base was replaced by a four-bp sequence in other lines. However, as no three-base duplication, such as the *Tpn1* transposon footprint, was observed in the four-bp sequence, it is challenging to speculate on the cause of this polymorphism.

To investigate the distribution of the newly identified transposons, a BLASTn search against *I. nil* sequences was performed. Two copies of *Tpn7-like* elements were found in the TKS genome sequence, which only differed in the length of the polythymine stretches (Supplementary Table S3). *Tpn20* was highly similar to *Tpn13*, which was inserted in the *EFP* gene (Morita et al. 2014), as well as to five transposon copies in the TKS genome, but exhibited multiple polymorphisms, including a single nucleotide substitution, as well as differences in the length of homopolymer sequences (Supplementary Table S4). No closely related transposons were found for *Tpn11*.

The same 1-bp deletion in intron 1, identical to the long-amplicon sequence in TKS, was found in the sequences of AK007, AK010, AK127, and AK205 (Fig. 3A). However, this deletion was not observed when it was analyzed using Sanger sequencing, which suggests that it was an error in the amplicon sequencing (Supplementary Fig. S2B). A comparison of the *Tpn11* sequences in AK010 and AK127 revealed differences in the length of the polythymine and polyadenine sequences (Supplementary Fig. S3). Although Sanger sequencing was attempted in these sequences, overlapping peaks prevented the determination of the exact length of the polyadenine stretch around 30 bp (Supplementary Fig. S3B). This was presumed to be due to PCR error, which resulted in polyadenine sequences of varying lengths being mixed within the amplicons. These results suggest that the length of homopolymer sequences may not always be accurately determined because of PCR and sequencing errors. However, some homopolymer sequences are consistent with reference sequences and can be accurately analyzed, which indicates that not all such sequences are problematic in determining length (Supplementary Fig. S1). In addition, the sequences obtained from lines other than TKS revealed deletions or duplications at the primer sites on both ends (Supplementary Fig. S4). These sequences could be corrected using the primer sequences and posed no practical issues.

In this study, long amplicons greater than 10 kb were successfully sequenced by circularizing the amplicons with T4 ligase and using a commercial plasmid DNA full-length analysis service. This method enabled the characterization of SRRs in *Tpn1* transposons, which are challenging to sequence using Sanger sequencing. Unlike primer walking with Sanger sequencers, this approach does not require primers or assembling sequences, thus saving time, labor, and cost. Although long-read amplicon sequencing services have recently been introduced in Japan, the cost of plasmid DNA full-length analysis remains less than one-tenth of those services. Further cost reductions are possible by preparing multiplex libraries and using in-house long-read sequencers; however, this requires an initial investment. While commercial long-read amplicon sequencing services, which are cost-competitive compared to plasmid DNA full-length analysis, are currently available in the United States, until similar services become widely available, the method described in this study remains the most cost-effective option for determining the full-length sequence of long PCR products.

## Materials and Methods

The plants used in this study are listed in Table 1, and their flower phenotypes are presented in Fig. 2. AK010 and AK127, which bear white or variegated flowers, were independently isolated from seeds of the commercial Violet line (Marutane Co., Ltd., Japan). AK205, which exhibits white or variegated flowers, was isolated from the progeny of a germinal revertant with fully colored flowers, derived from Q1072. AK007 and Q1072 are sublines of line K37. Genomic DNA was extracted using either the Genomic-tip 500/G (QIAGEN, Germany) or the GENE PREP STAR PI-480 (KURABO, Japan), as previously described (Hoshino et al. 2016). The 4–11 kb fragments of the *DFR-B* sequence were amplified by PCR using the primers DP-LF (5′-TTAACATGAGGGGATTGCATGTCACTTTCA-3′) and D3U-LR (5′-CATAAATCTGGTTCGAGTGGCAATCTAACT-3′) with Ex-Premier DNA Polymerase (TAKARA, Japan). The PCR conditions were as follows: 94°C for 1 min, followed by 35 cycles of 98°C for 10 s, 57°C for 15 s, and 68°C for 7 min. The PCR product obtained was precipitated with ethanol and dissolved in 17 μL of Milli-Q water. Subsequently, 2 μL of T4 Ligase buffer (New England Biolabs, USA) and 1 μL of T4 Polynucleotide Kinase (10 U/μL; TAKARA) were added, and the reaction was incubated at 37°C for 30 min to phosphorylate the 5’ ends. An additional 1 μL of T4 Ligase (200 U/μL; New England Biolabs) was then added, and the mixture was incubated at room temperature for 2 h to circularize the DNA. The circularized DNA was recovered by ethanol precipitation, dissolved in low TE buffer pH 8.0 (10 mM Tris and 0.1 mM EDTA), and a 10 μL solution with a concentration of 50 ng/μL was prepared. Nanopore sequencing was performed using the Plasmid-EZ service, which provides full plasmid DNA sequence analysis (AZENTA, Japan). For sequences suspected to be sequencing errors, Sanger sequencing was performed using the DNA that was used as the template for the amplicon sequence. Sequences were analyzed using SnapGene version 5.0.8 (GSL Biotech LLC, USA) and ATGC-MAC version 7.2.1 (NIHON SERVER, Japan).

**Table 1.**
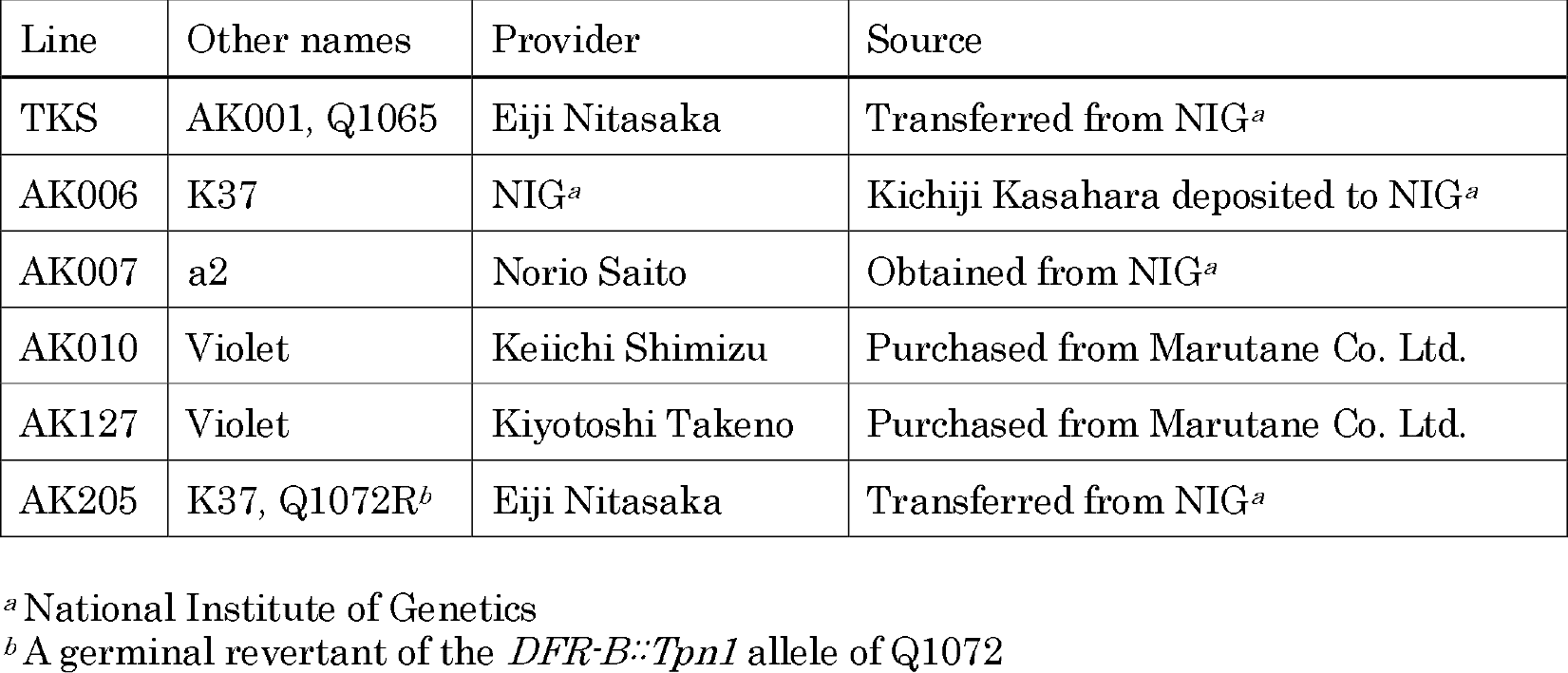
Plant lines used this study.

The raw sequencing data have been deposited in the DNA Data Bank of Japan (DDBJ)/BioProject under accession number PRJDB18807.

## Acknowledgments

The authors thank Eiji Nitasaka, Keiichi Shimizu, Kiyotoshi Takeno, and the late Norio Saito for providing the *I. nil* seeds. Moreover, thanks are extended to Kazuyo Ito, Tomoyo Takeuchi, Kiyoko Kuzunishi, and Naoko Koyama for their technical assistance, along with the Model Organisms Facility and Trans-Omics Facility, NIBB Trans-Scale Biology Center. JSPS KAKENHI supported part of this study with grants to AH (21K06239) and SN (23KJ1004).

