## Supplementarly Figures and Tables for "Labor- and cost-effective long-read amplicon sequencing using a plasmid analysis service: Application to transposon-inserted alleles in Japanese morning glory"

**Supplementary Table S1.** Summary statistics of read lengths

| Line | Minimum | Mean | Max | Q1 [a] | Median | Q3 [a] | N50 |
|---|---|---|---|---|---|---|---|
| TKS | 88 | 1930.1 | 19620 | 1173.5 | 1762 | 2545.5 | 2379 |
| AK006 | 119 | 2390.83 | 23,610 | 861.75 | 1,643 | 3125.25 | 3,767 |
| AK007 | 114 | 2318.5 | 90,511 | 589 | 1,387 | 3088.5 | 4,296 |
| AK010 | 123 | 2979.29 | 14,988 | 1044 | 2,057 | 4247 | 5,042 |
| AK127 | 136 | 2892.11 | 19,674 | 1106.5 | 2,123 | 4022.75 | 4,527 |
| AK205 | 111 | 2773.83 | 12,741 | 992.5 | 2,205 | 3743.5 | 4,183 |

[a] Read length at the interquartile range corresponds to Q1 and Q3, representing the 25th and 75th percentiles of read length, respectively.

**Supplementary Table S2.** Quality assessment of assembly

| Line | Length [a] | Reads | Mapped reads | Unmapped reads | Supplementary [b] |
|---|---|---|---|---|---|
| TKS | 4,130 | 1,608 | 1,585 (98.56%) | 23 (1.43%) | 43 (2.67%) |
| AK006 | 10,544 | 1,366 | 1,124 (82.28%) | 242 (17.71%) | 48 (3.51%) |
| AK007 | 10,930 | 633 | 436 (68.87%) | 197 (31.12%) | 16 (2.52%) |
| AK010 | 11,335 | 937 | 876 (93.48%) | 61 (6.51%) | 30 (3.2%) |
| AK127 | 11,336 | 775 | 740 (95.48%) | 35 (4.51%) | 20 (2.58%) |
| AK205 | 10,447 | 567 | 542 (95.59%) | 25 (4.4%) | 88 (15.52%) |

[a] Length of the assembled sequences.

[b] Reads that span the 3´–5´ boundary of the linearized contig sequence.

**Supplementary Table S3.** *Tpn7* related elements

| Transposon | Line | Scaffold [a] | Start | End | Length | Identity (%) | Strand |
|---|---|---|---|---|---|---|---|
| *Tpn7* | AK007 | | | | 6,776 | | + |
| *Tpn7-like* | TKS | 3415 | 502,138 | 508,930 | 6,793 | 99.75 | − |
| *Tpn7-like* | TKS | 3047 | 597,586 | 604,379 | 6,794 | 99.74 | + |

[a] Scaffold number of the Asagao_1.1 genome (GCF_001879475.1).

**Supplementary Table S4.** *Tpn20* related elements

| Transposon | Line | Scaffold [a] | Start | End | Length | Identity (%) | Strand |
|---|---|---|---|---|---|---|---|
| *Tpn20* | AK205 | | | | 6,258 | | − |
| *Tpn13* | AK028 | | | | 6,284 | 99.55 | + |
| *Tpn20-like* | TKS | 0782 | 811,066 | 817,335 | 6,270 | 99.79 | + |
| *Tpn20-like* | TKS | 0900 | 89,223 | 95,490 | 6,268 | 99.82 | − |
| *Tpn20-like* | TKS | 1074 | 4,279,704 | 4,285,964 | 6,267 | 99.74 | + |
| *Tpn20-like* | TKS | 2833 | 2,030,717 | 2,036,978 | 6,267 | 99.76 | − |
| *Tpn20-like* | TKS | 1721 | 616,967 | 623,240 | 6,276 | 99.65 | − |

[a] Scaffold number of the Asagao_1.1 genome (GCF_001879475.1).

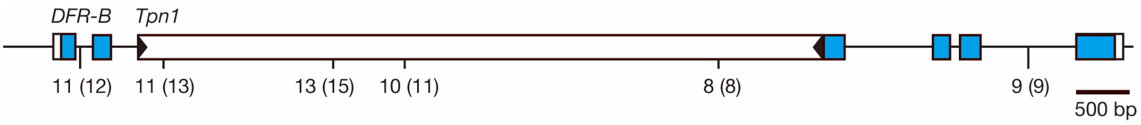


**Supplementary Figure S1.** Putative sequence errors in the polythymine stretch of *DFR-B::Tpn1* in AK006 resulted in a few base pair deletions. The boxes represent exons, with the coding regions shaded in blue. The box with black arrowheads at the ends represents *Tpn1*. Vertical bars indicate the positions of polythymine stretches of 8 bp or longer. The numbers below the bars indicate the polythymine lengths (bp) in the long-read amplicon sequence of AK006, with those of the reference genome sequence (GCF_001879475.1) and *Tpn1* sequence (D37795) shown in parentheses.

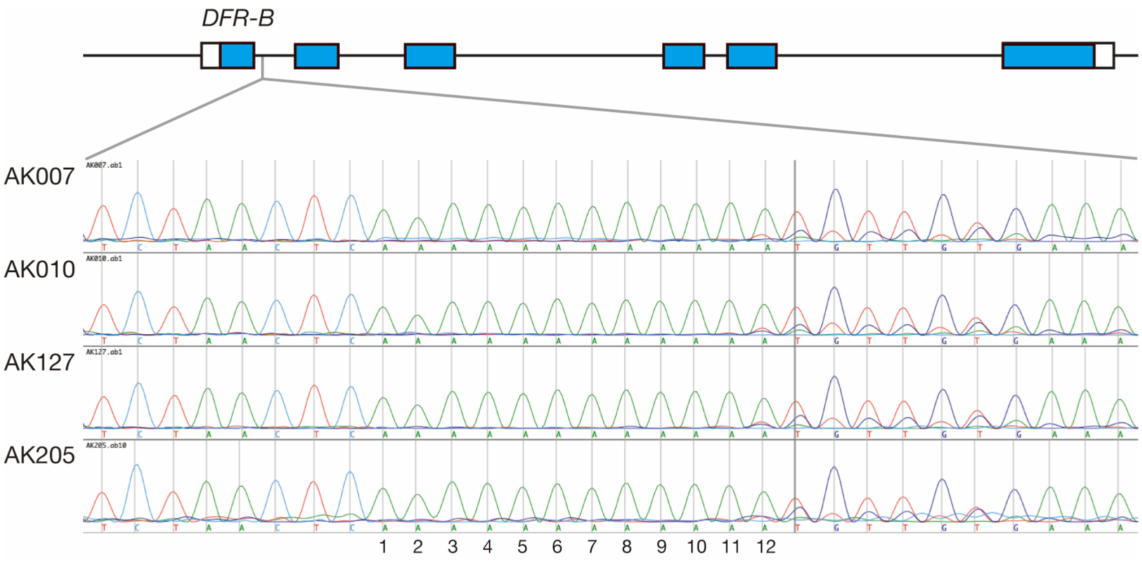


**Supplementary Figure S2.** Sanger sequencing reveals a sequence error in the 12-bp polythymine stretch analyzed by the DNA sequencing of the full-length plasmid. A schematic drawing of the *DFR-B* gene is shown at the top, with boxes representing exons and the coding regions shaded in blue. The Sanger sequence electropherograms are displayed at the bottom. The polythymine stretch was sequenced using the reverse primer DFR-RV6 (5´-GGGTCCTTGGAATCGAAATC-3´). While the assembled Nanopore reads show an 11-bp polythymine stretch, Sanger sequencing reveals the actual sequence is a 12-bp stretch, which matches the reference genome sequence (GCF_001879475.1). The sequence after the 12th adenine shows overlapping peaks, suggesting the presence of homopolymers of varying lengths generated by PCR.

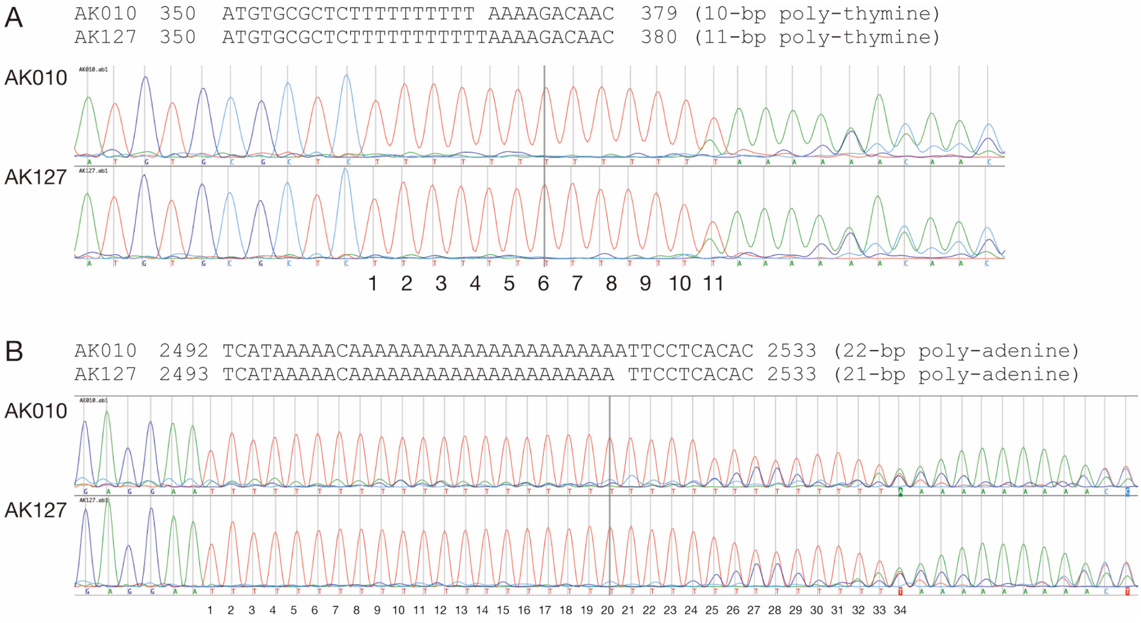


**Supplementary Figure S3.** Apparent length polymorphism of *Tpn11* between AK010 and AK127. (A) A polythymine stretch begins at position 350 of *Tpn11*. Below, the Sanger sequencing electropherogram using the forward primer DFR-Fw1 (5′-ATATGCCAAGAAAATGACTGGAT-3′) is shown. (B) A polyadenine stretch begins at positions 2492 and 2493 of *Tpn11* in AK010 and AK127, respectively. Below, the Sanger sequencing electropherogram using the reverse primer Tpn11-Rv1 (5′-TACGCACTGTTGGAGGTTGG-3′) is presented. Because of overlapping peaks following the homopolymers, particularly in the longer polyadenine stretch, the exact lengths of the homopolymers are inaccurate in both long-amplicon and Sanger sequencing.

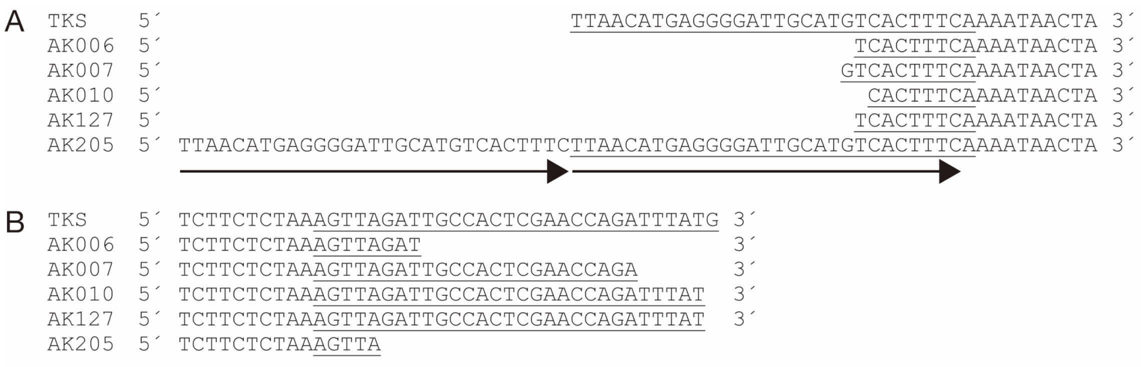


**Supplementary Figure S4.** Rearrangement at both ends of the fragment sequences: 5´ end (A) and 3´ end (B) sequences. The primer sequences are underlined, and an artificial tandem duplication at the 5´ end is indicated by the arrows.
